# Aperiodic neural timescales in prefrontal cortex dilate with increased task abstraction

**DOI:** 10.1101/2025.04.21.649913

**Authors:** Dillan Cellier, Justin Riddle, Ryan Hammonds, Flavio Frohlich, Bradley Voytek

## Abstract

Navigating everyday environments requires that the brain perform information processing at multiple different timescales. For example, while watching a movie we use sensory information from every video frame to construct the current movie scene, which itself is continuously integrated into the narrative arc of the film. This critical function is supported by sensory inputs propagating from dynamic sensory cortices to association cortices, where neural activity remains more stable over time. The hierarchical organization of cortex is therefore reflected in a gradient of neural timescales. While this propagation of inputs up the cortical hierarchy is facilitated by both rhythmic (oscillatory) and non-rhythmic (aperiodic) neural activity, traditional measures of oscillations are often confounded by the influence of aperiodic signals. The reverse is also true: traditional measures of aperiodic neural timescales are influenced by oscillations. This makes it difficult to distinguish between oscillatory and timescale effects in cognition. Here, we analyzed electroencephalography (EEG) data from participants performing a cognitive control task that manipulated the amount of task-relevant contextual information, called task abstraction. Critically, we separated aperiodic neural timescales from the confounding influence of oscillatory power. We hypothesized that neural timescales would increase during the task, and more so in high-abstraction conditions. We found that task abstraction dilated the aperiodic neural timescale, as estimated from the autocorrelation function, over prefrontal cortical regions. Our findings suggests that neural timescales are a dynamic feature of the cerebral cortex that change to meet task demands.

**Significance Statement:** Though often thought of as indexing anatomical brain structure, neural timescales have recently come under the spotlight for their variability across different task conditions. Here, we investigated whether neural timescales dynamically change during cognitive tasks that require a greater level of contextual control. Previous work quantified neural timescales using the autocorrelation function, but this approach conflates oscillatory and aperiodic contributions to the timescale. Thus, we decomposed the neural timescale into oscillatory and aperiodic components, and found that aperiodic-derived timescale measures dynamically change with contextual control. Oscillatory power also exhibits change in response to task demands, reaffirming previous findings. These results together suggest that both established oscillatory and novel aperiodic measures of neural timescales can dynamically change to meet task demands.

## Introduction

Neural activity evolves over space and time; while two spatially proximate cortical regions are generally similar, the signals of two spatially *distant* regions are generally dissimilar (i.e., spatial autocorrelation)(1). Temporal autocorrelation— correlating a neural signal with a time-lagged version of itself— reveals that association cortices are dominated by slower neural timescales than sensory cortices. This results in a temporal hierarchy of neural timescales that mirrors the hierarchical arrangement of the brain (2–12). Neural timescales were conceptualized as *intrinsic* neural timescales—that is, an emergent property of the anatomical network structure of the brain (3), however, new empirical evidence calls into question whether neural timescales reflect a static network property of the brain, or alternatively, reflect the temporal bounds within which a brain region can flexibly respond to task demands (13). Previous work suffered from two primary confounds that are addressed by the current investigation.

First, previous work did not utilize tasks that isolate shorter-versus-longer cognitive timescales by manipulating the integration of information required to complete the task— known as task abstraction. Task abstraction requires neural integration of lower-order inputs in higher-order cortical regions (5,14,15), and neural timescales are thought to reflect this integration (3,16–18). The regions of cortex with the longest timescales are also the regions most closely associated with greater task abstraction (5,12,19,20). The second confound of previous work is the simple sampling from the autocorrelation function (ACF) to estimate neural timescales. This metric conflates aperiodic and oscillatory signals. That is, the observation of slower timescales from the ACF of one condition compared to another could be by virtue of differences in the power of low-frequency neural oscillations, or a difference in the proportions of low-frequency aperiodic activity. This may explain why recent studies examining task-related changes in neural timescales report conflicting findings (13,16,21–23). While some studies suggest that task-state timescales are longer than resting or pre-trial timescales (24,25), others conclude that timescales are task invariant (26).

To address these confounds, we quantified neural timescales from scalp EEG while human participants performed a task designed to drive abstraction. We hypothesized that high-abstraction conditions would dilate neural timescales. To test this, we re-analyzed two independent EEG datasets from Riddle et al., 2020 and Riddle et al., 2021, where abstraction was operationalized as a decision tree of possible actions, organized by a superordinate set of context rules (29). Critically, the task employed in these studies isolates the effects of task abstraction from those of task difficulty, an often overlooked confounding factor in cognitive tasks. We quantified neural timescales using traditional metrics, as well as modeling the aperiodic component and oscillatory power as they changed from a pre-trial baseline to stimulus processing. We found that traditionally measured neural timescales increase with stimulus onset in prefrontal and posterior channels. While prefrontal aperiodic timescales exhibited selective lengthening in response to abstraction, prefrontal theta oscillatory power increased with difficulty, and posterior alpha oscillatory power showed mixed selectivity. These findings suggest that aperiodic neural timescales are not wholly intrinsic, but in fact exhibit condition-specific dynamism that may reflect the demands of integration of a given task state.

## Methods

### Novel separation of neural timescales into aperiodic and oscillatory components

Electrophysiological neural signals are products both of oscillatory and non-oscillatory, or aperiodic, activity (30). However, studies of neural timescales typically do not disambiguate the contributions of these distinct types of signals to the timescale— traditional metrics that sample from the raw autocorrelation function (ACF) instead blend the aperiodic components and oscillatory components of the signal (see **Figure S1E**). Here, we rely on two types of transformations of the timeseries data to disentangle aperiodic and oscillatory changes during a task: the ACF, and the power spectral density (PSD) (**Figure S1B and C**), which are mathematically related to each other according to the Wiener-Khinchin theorem (24,31–33).

For a timeseries 𝑋, the autocorrelation function is calculated as:

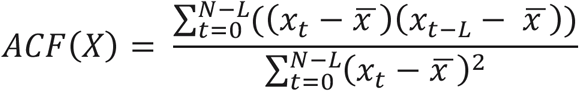

where 𝑁 is the length of the signal, 𝐿 is the lag, and 𝑡 is the sample at every point from 𝑡 = 0 to 𝑁 − 𝐿. Moreover, the ACF of purely aperiodic activity may be modeled as an exponential decay function described by 𝑒^−𝑡/𝜏^, with decay constant 𝜏. This can be used to derive the number of lags that it takes for Pearson’s correlation values to drop to a value of 𝑦. The decay constant can be approximated by:

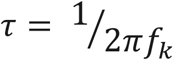

where 𝑓_𝑘_ represents the knee frequency of the power spectrum. The knee frequency (34), or the frequency value at which the power values go from being roughly constant to power-law distributed (i.e., the frequency at which the power spectrum “bends”), is a part of the Lorentzian function (and spectral approximation of the exponential decay ACF):

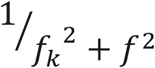

used to describe the PSD of aperiodic activity (24). Timeseries composed of slow aperiodic signals possess higher tau, 𝜏, values in the ACF and lower knee frequencies, 𝑓_𝑘_, in PSD space when compared to timeseries composed of faster signals (**Figure S1A**). Thus, tau reflects the longer “memory” of slower signals (it takes more lags to “forget”) (**Figure S1B**). Low PSD knees of slow signals reflect greater power for low-frequencies relative to high (**Figure S1C**). Highly stochastic signals that are aperiodic and “fast,” with little temporal autocorrelation, yield virtually flat ACFs and PSDs—reflecting limited memory and equal power across both low and high frequencies.

The timescales of purely aperiodic activity, therefore, are reflected in the tau of the ACF and the knee of the PSD. Similarly, the frequency and power of purely oscillatory signals are reflected in their ACFs and PSDs. In the ACF, neural oscillations are found as rhythmic peaks and troughs in the autocorrelation that reflect the frequency of the oscillation (**Figure S1E**). In the PSD, oscillations appear as Gaussian “peaks” at the center frequency of the oscillation above the backdrop of the aperiodic signal (**Figure S1F**). Neural oscillations are therefore also captured in previous findings of neural timescales in cortex, but should be separated from aperiodic activity.

One common approach for measuring timescales from the raw ACF is to set a threshold on the y-axis (representing the correlation values) and extract the point on the x-axis (the lag number) at which the ACF drops past this threshold. This threshold can be set to the 50% correlation mark (12), thus this metric is referred to here as the 50-Crossing value (**Figure 1C**). These N-Crossing values will be greater for slower timescales, and lesser for faster timescales (**Figure 1C insert**). Often, the ACF is mirrored over the x-axis, and the width of the function measured at the y-axis threshold is taken as the timescale, i.e., the autocorrelation window approach (17,33).

**Figure 1:**
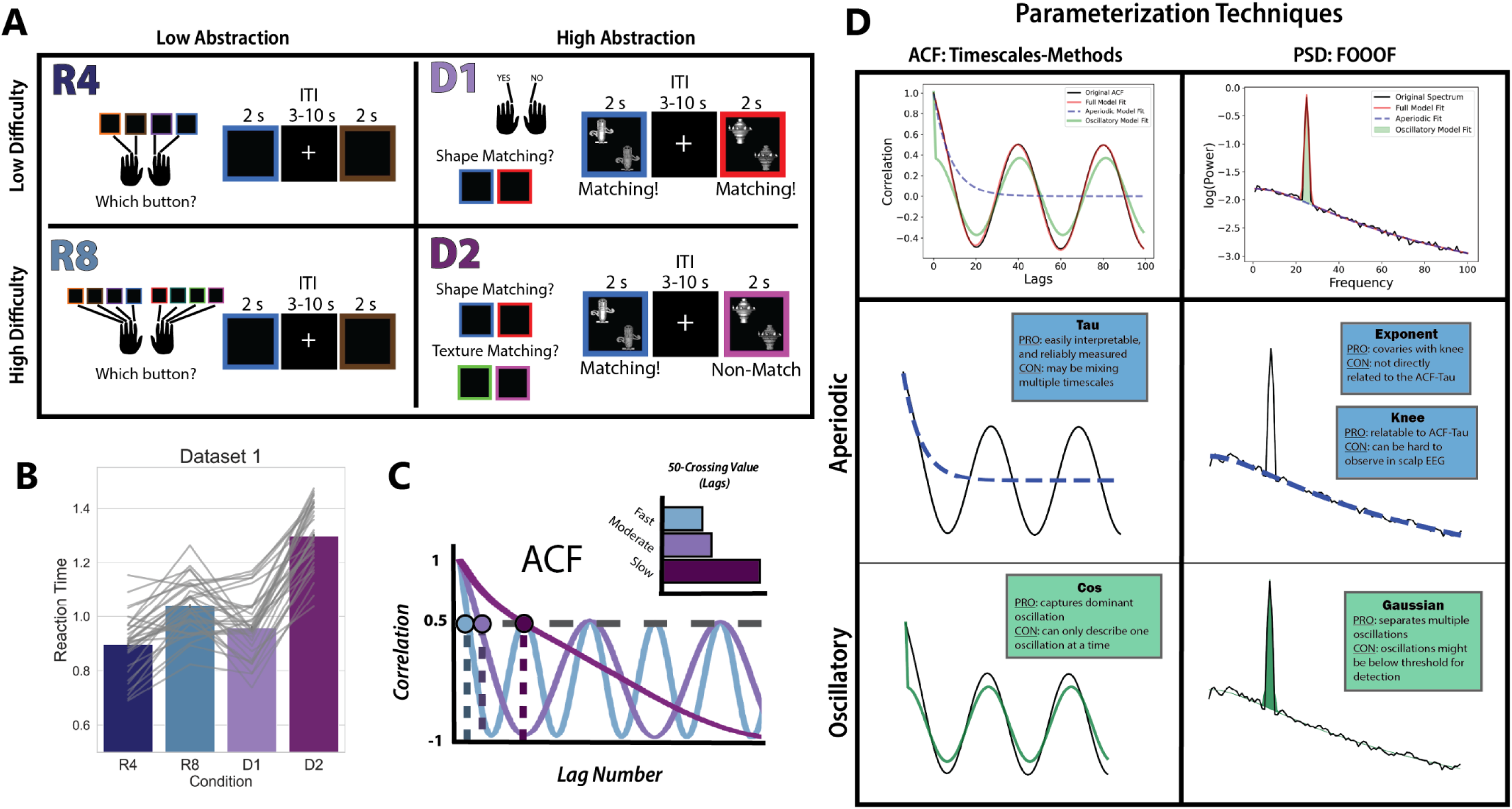
Task Schematic and EEG Analyses. A) Task schematic. This block-design task included 4 conditions that varied along two dimensions: difficulty and abstraction. In the R4 condition (low difficulty, low abstraction) participants had to respond with a button press corresponding to the color they observed on each trial. This required participants to keep in mind a set of 4 stimulus-response mappings throughout the course of the block. The R8 condition (high difficulty, low abstraction) had the same design as R4, except that participants needed to hold 8 stimulus-response mappings in mind over the course of the block. In the D1 condition (low difficulty, high abstraction), participants were making a match-nonmatch judgement on every trial. The two stimuli they were evaluating could vary in both shape and texture. Which feature was relevant for the match-nonmatch judgment was cued by the color of the square surrounding the items. In the D1 condition, the cued color corresponded to the same feature for the entirety of the block. However, in the D2 condition (high difficulty, high abstraction), the cued feature could change trial-to-trial. This meant that on every trial, the participant needed to recall which feature the color of the square cued, evaluate whether the stimuli matched on that feature dimension, and press a button for “match” or “nonmatch.” B) Barplots of the participant-averaged reaction times for each condition, with individual reaction times plotted in grey on top. The high difficulty, high abstraction condition D2 had the longest reaction time in both datasets. C) One common method for estimating neural timescales entails calculating the ACF of a timeseries, and then identifying the lag (x-axis value) at which the ACF crosses a given correlation threshold, N (in this example, N=0.50). This gives a relative measure between timeseries, illustrated in E, where slower signals have larger N-crossing values. This metric will correlate well with the decay rate of the ACF, but mixes the contributions of aperiodic and oscillatory activity. D) An overview of the two procedures used to parameterize the ACF and PSD. Each toolbox (Timescales-methods and FOOOF) produces both aperiodic and oscillatory parameters. The Timescales-methods toolbox for ACF parameterization derives a Tau parameter, which captures the decay rate of the aperiodic component of the ACF. It can also model the oscillatory component of the ACF using a damped cosine function, producing the parameter Cos. The FOOOF toolbox for PSD parameterization produces several parameters related to the aperiodic component of the signal, among them the knee parameter and the exponent parameter. Though the knee parameter most directly relates to the ACF-Tau, it is not reliably visible in real neural data—especially that which is derived from non-invasive imaging. The exponent parameter, which describes the slope of the aperiodic signal, covaries with the knee.

### Modeling the Neural Timescale using Power Spectral Density

We quantified oscillatory power and aperiodic timescales using PSD parameterization (**Figure 1D**). The PSD of each trial and time window was calculated using six-cycle Morlet wavelets, for frequencies between 2 and 50 Hz, and on mirror-padded timeseries. Mirror padding consists of appending a reflected version of the timeseries before and after each signal segment, before transformation to the frequency domain, and was conducted to add length to the signal and improve frequency resolution(35). With a one-second time window for analysis, our lower bound was 2 Hz. We model PSDs using the Python-based, open-source FOOOF toolbox (36). Briefly, the FOOOF toolbox estimates the aperiodic signal of a PSD by deriving a rough first estimate of the line of best fit, iteratively fitting and removing Gaussian peaks over putative oscillatory peaks, and re-estimating the aperiodic signal from the non-oscillatory residuals of the spectrum. The FOOOF settings used for model fitting were as follows: peak width limits were set to a minimum of 2 and a max of 12, the maximum number of peaks allowable were 4, the peak threshold was set to 2.5, and the aperiodic mode was fixed.

From each model, the estimated oscillatory Gaussian peak parameters (amplitude, bandwidth, and center frequency) in the theta (4-8 Hz), and alpha (8-12 Hz) ranges were saved for later analysis. We excluded delta oscillations from analysis due to their being at the lower bound of our frequency range (2 Hz), and therefore unreliable to model. We additionally documented the aperiodic-corrected bandpower in each of these oscillatory band ranges. That is, we calculated the average of the raw power values between the low and high frequency cut-off of the band range, minus the average power of the aperiodic fit between those same frequencies (*PSD-Theta* and *PSD-Alpha*). This was necessary because, often, oscillatory peaks were not successfully detected by FOOOF, resulting in missing oscillatory estimates for certain subsets of the data. We therefore had two measures of oscillatory activity: Gaussian parameters, which give a binary metric of whether oscillatory peaks are present or not, and aperiodic-corrected raw bandpower, which gives a continuous measure of power (and near-zero power when an oscillation is not present).

Lastly, given that the knee parameter of the PSD is the most directly mathematically relatable to the tau parameter of the ACF (24), it would be ideal to measure the knee parameter from our scalp EEG. However, the aperiodic knee was not uniformly present across raw PSDs; this is not uncommon to observe in EEG data, and particularly in cases where the time window of data acquisition is insufficiently long to model the true knee frequencies. We therefore analyzed the aperiodic exponent (*PSD-Exponent*) as a covariate of aperiodic knee frequency changes that would be introduced by changes in neural timescales. We offer a supplementary analysis of the PSD data, fit with a knee parameter, in order to investigate the validity of excluding the knee parameter from our first set of analyses (**Supplementary Notebook 00**).

### Modeling the Aperiodic Neural Timescale using the Autocorrelation Function

The metric of interest derived from modeling of the ACF was the tau parameter (*ACF-Tau*), which reflects the aperiodic component of the ACF. We also calculated the traditional 50-Crossing metric (labeled as *50-Crossing*) from the raw ACF, which conflates oscillatory and aperiodic contributions to the timescale. Here, the tau parameter provides a novel approach to estimating aperiodic temporal autocorrelation that might more directly reflect the neural timescale compared to the exponent of the aperiodic signal from PSD.

For each timeseries from the pre- and post-stimulus time windows, for every trial, and for each set of averaged electrodes, we calculate an ACF using the statsmodels Python package. We then modeled ACFs using an in-house, Python-based toolbox named Timescales-Methods, openly available at https://voyteklab.com/timescale-methods/. This toolbox operates by estimating the timescale of an exponential decay function, which models the aperiodic contribution to the ACF, given by the formula:

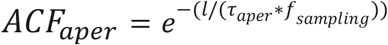

where 𝑙 is an array of lag values of length <= the length of the signal, 𝜏*_aper_* is the timescale, and 𝑓*_sampling_* is the sampling rate of the signal. The tau parameter, 𝜏*_aper_*, was then transformed to be more comparable to the raw ACF 50-Crossing metric. This was done by calculating the predicted lag at which the ACF crosses the 50% correlation threshold, given tau, via the formula:

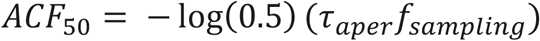

The oscillatory component of the ACF is modeled here as a damped cosine function, given by the formula:

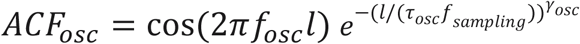

However, the parameterization of oscillatory effects on the ACF proves challenging. Multiple oscillations appear as overlapping rhythms in the ACF, thus, the PSD is superior for separating distinct oscillatory rhythms from one another. We ultimately exclude the ACF damped cosine parameters from our main analyses due to our modeling procedure only capturing the most dominant oscillation, though perhaps future ACF modeling procedures could be developed to separate multiple oscillations from the ACF. We offer a supplementary analysis of the oscillatory parameters extracted by our ACF modeling in **Supplementary Notebook 05**. Our main oscillatory findings focus on estimates of oscillatory activity from the PSD, instead of from the ACF.

We experimented with model-fitting the raw ACF with and without the oscillatory component (damped cosine function) as a matter of hyperparameter tuning. The rationale for testing this hyperparameter choice is that the damped cosine includes an exponential decay term, which may impair estimation of the true decay rate of the aperiodic component of the ACF (especially for timeseries with high-powered oscillations). The main ACF-Tau findings reported in the Results section and shown in **Figure 4** are from models with the oscillatory component *excluded*, while results from models with the oscillatory component *included* are in **Supplementary Notebook 04 and 05**. The statistical results of both procedures are reported below in **Table 1**. We additionally include a visualization of the relationship between ACF-Tau parameters and PSD-derived oscillatory bandpower, with the goal of ensuring that our ACF-Tau results are not driven specifically by oscillatory power that the modeling failed to remove (**Supplementary Notebook 06**).

**Table 1:** Results from a One-Way ANOVA of all conditions, and a Two-Way ANOVA of Abstraction and Difficulty, calculated for each neural metric and each region of interest. Green highlights indicate findings for which p < 0.05.

|  |  |  | One Way ANOVA |  |  | Two Way ANOVA |  |  |  |  |  |  |  |  |
| --- | --- | --- | --- | --- | --- | --- | --- | --- | --- | --- | --- | --- | --- | --- |
|  |  |  | Main Effect of Condition |  |  | Main Effect of Abstraction |  |  | Main Effect of Difficulty |  |  | Interaction |  |  |
|  |  |  | F-Statistic | dof | P-Value | F-Statistic | dof | P-Value | F-Statistic | dof | P-Value | F-Statistic | dof | P-Value |
| Dataset: | Channel Cluster: | Metric: |  |  |  |  |  |  |  |  |  |  |  |  |
| Dataset 1 | Frontal | ACF Tau (fit without osc) | 4.40 | 30 | 0.006 | 9.36 | 30 | 0.005 | 0.50 | 30 | 0.483 | 4.69 | 30 | 0.039 |
|  |  | ACF Tau (fit with osc) | 8.53 | 30 | 0.000 | 13.73 | 30 | 0.001 | 2.76 | 30 | 0.107 | 7.99 | 30 | 0.008 |
|  |  | ACF 50-Crossing | 37.13 | 30 | 0.000 | 46.11 | 30 | 0.000 | 31.78 | 30 | 0.000 | 25.21 | 30 | 0.000 |
|  |  | Theta Bandpower | 5.46 | 30 | 0.002 | 4.65 | 30 | 0.039 | 12.00 | 30 | 0.002 | 0.82 | 30 | 0.373 |
|  |  | PSD Exponent | 2.37 | 30 | 0.076 | 2.03 | 30 | 0.165 | 3.88 | 30 | 0.058 | 1.43 | 30 | 0.241 |
|  | Posterior | ACF Tau (fit without osc) | 0.78 | 30 | 0.509 | 0.90 | 30 | 0.350 | 0.87 | 30 | 0.360 | 0.41 | 30 | 0.525 |
|  |  | ACF Tau (fit with osc) | 0.84 | 30 | 0.477 | 2.25 | 30 | 0.144 | 0.08 | 30 | 0.783 | 0.00 | 30 | 0.962 |
|  |  | ACF 50-Crossing | 3.12 | 30 | 0.030 | 2.57 | 30 | 0.119 | 5.03 | 30 | 0.032 | 0.27 | 30 | 0.608 |
|  |  | Alpha Bandpower | 19.14 | 30 | 0.000 | 28.30 | 30 | 0.000 | 24.64 | 30 | 0.000 | 5.08 | 30 | 0.032 |
|  |  | PSD Exponent | 0.61 | 30 | 0.609 | 0.58 | 30 | 0.451 | 1.19 | 30 | 0.284 | 0.22 | 30 | 0.641 |
| Dataset 2 | Frontal | ACF Tau (fit without osc) | 4.80 | 22 | 0.004 | 7.47 | 22 | 0.012 | 2.76 | 22 | 0.111 | 3.24 | 22 | 0.086 |
|  |  | ACF Tau (fit with osc) | 1.82 | 22 | 0.152 | 2.58 | 22 | 0.122 | 1.28 | 22 | 0.270 | 1.67 | 22 | 0.210 |
|  |  | ACF 50-Crossing | 19.77 | 22 | 0.000 | 38.04 | 22 | 0.000 | 15.05 | 22 | 0.001 | 3.77 | 22 | 0.065 |
|  |  | Theta Bandpower | 3.79 | 22 | 0.014 | 2.78 | 22 | 0.110 | 9.19 | 22 | 0.006 | 1.24 | 22 | 0.278 |
|  |  | PSD Exponent | 1.54 | 22 | 0.213 | 0.37 | 22 | 0.548 | 3.49 | 22 | 0.075 | 0.60 | 22 | 0.448 |
|  | Posterior | ACF Tau (fit without osc) | 0.35 | 22 | 0.789 | 0.07 | 22 | 0.800 | 0.02 | 22 | 0.895 | 1.94 | 22 | 0.177 |
|  |  | ACF Tau (fit with osc) | 0.68 | 22 | 0.569 | 0.00 | 22 | 0.994 | 0.35 | 22 | 0.560 | 3.79 | 22 | 0.064 |
|  |  | ACF 50-Crossing | 0.82 | 22 | 0.485 | 0.05 | 22 | 0.834 | 3.56 | 22 | 0.073 | 0.09 | 22 | 0.765 |
|  |  | Alpha Bandpower | 26.09 | 22 | 0.000 | 50.40 | 22 | 0.000 | 29.16 | 22 | 0.000 | 4.97 | 22 | 0.036 |
|  |  | PSD Exponent | 1.01 | 22 | 0.395 | 2.27 | 22 | 0.146 | 0.09 | 22 | 0.762 | 0.40 | 22 | 0.533 |

Our 50-Crossing metric was sampled from the raw ACFs, with linear interpolation of the two lag values between which the correlation dropped below 50. We additionally sampled the raw ACF at the slightly lower 0-Crossing point, given debate over which metric is best (25). The results of this alternative measure can be found in **Supplementary Notebook 03**.

Lastly, we visualize the results of our ACF-Tau, PSD-Exponent, and ACF 50-Crossing measures in relation to event-related changes in voltage pre-to-post-stimulus **Supplementary Notebook 07**. We do not find compelling evidence that the results presented here are primarily driven by event-related potentials.

### Experimental Task and Behavior

The hierarchical cognitive control task developed for Riddle et al., 2020 was the same used for Riddle et al., 2021. Critically, this task manipulates task abstraction separately from task difficulty in a 2-by-2 design. This strategically enabled us to separate any observed effects of task abstraction on neural timescales from the effect of task difficulty or overall response time. The task used a blocked design, with two levels of task difficulty and two levels of task abstraction **(Figure 1A)**. Here, task abstraction is operationalized as the number of context-dependent rules to be parsed to determine the correct motor output, and task difficulty as the number of rules memorized in each condition.

The response tasks (low-abstraction) consisted of two conditions and required only stimulus-to-response mapping— there was no context-dependent stimulus evaluation. The response tasks required either four (R4) or eight (R8) stimulus-to-response mapping rules. That is, participants observed a colored square for up to two seconds and were tasked with pressing the one button of four (or eight) corresponding to that color. While the R4 condition is low-difficulty *and* low-abstraction, the R8 condition is low-abstraction but high-difficulty. The dimension tasks, D1 and D2, required the integration of contextual information (high abstraction). The dimension tasks required a match-nonmatch judgment of two stimuli, where the stimuli could vary along two dimensions: shape and texture. The dimension along which the match-nonmatch judgment was made was determined by the color of the square surrounding the items. In the low difficulty/high abstraction condition (D1), there was a single dimension indicated by the color of the surrounding square (shape and texture blocks were counterbalanced and order randomized). In the high difficulty/high abstraction block (D2), there were two dimensions that could be indicated by the color of the square, and the relevant dimension could change trial-by-trial. The intertrial interval used for pre-stimulus baseline EEG data was jittered between 3 and 10 seconds, and the order of the conditions was randomized across blocks. Task rules were explicitly instructed prior to beginning the task, and participants had the opportunity to practice the task before data recording began. Detailed analysis of the behavioral data from these studies can be found in the original papers. Participants were slower and less accurate for the R8 condition compared to the D1 condition, and thus the difference between high abstraction and low abstraction conditions could not be explained by task difficulty **(Figure 1B)**.

### Preprocessing EEG Data

EEG data from both Dataset 1 (Riddle et al., 2020) and Dataset 2 (Riddle et al., 2021) contained recordings from 31 (18 females, mean age of 20) and 23 unique participants (19 females, mean age 19.7 years). The data were collected with approval from the University of California Berkeley Committee for Protection of Human Subjects and the University of North Carolina at Chapel Hill’s institutional review board. Written consent was obtained prior to data collection. These data were acquired with permission from the authors in preprocessed, epoched formats for the task data and raw, continuous formats for the resting-state data.

Details on the original preprocessing of Dataset 1 task data are reported in Riddle et al., 2020. The EEG data were acquired using a 64-channel active electrode BioSemi ActiveTwo amplifier, with electrodes organized according to the standardized 10-20 system. The data were filtered between 0.1-100 Hz using a two-way least-squares finite impulse response (FIR) filter. Data were epoched from -1000 to +2000 ms around stimulus onset before inspection for noisy epochs, rejection and interpolation of noisy channels, and rejection of eyeblinks using extended info-max independent component analysis (ICA)(37). No baseline correction was applied.

The raw, unprocessed resting-state data consisted of 5 minutes of eyes-open resting-state, continued by 5 minutes of eyes-closed resting-state, and were collected at the beginning of each experimental session (prior to task blocks). Due to relatively higher amounts of physiological noise in the resting-state session, we preprocessed this data in a series of steps that deviated somewhat from the protocol followed for task data. All preprocessing performed on these resting-state data were performed using MNE-Python version 1.61 (38,39). The data were first rereferenced to the average of channels P9 and P10 (Kappenman et al., 2021), bandpass filtered between 0.1-100 Hz using an overlap-add FIR filter, plotted, and inspected both for noisy electrode channels and noisy periods of time. We conducted ICA on the continuous data using the FastICA algorithm and 20 principal components (41). Independent components corresponding to eyeblink data were rejected. Data were then segmented into consecutive 1000 ms segments, with the goal of matching the length of pre- and post-stimulus windows of task data. These epoched data were then submitted to a regression of any ocular artifacts (Gratton et al., 1983). We visually inspected and rejected any noisy epochs that were not already identified during the previous annotation of bad data segments, and saved these data for later analysis.

Dataset 2 EEG data (Riddle et al., 2021) were recorded using a high-density 128-Channel electrode system from HydroCel Geodesic Sensor Net and EGI system. These data were collected as a baseline session for a multi-session study of the effects of transcranial alternating current stimulation (tACS) on EEG activity; the tACS related data is not analyzed here. The preprocessing of task data entailed filtering between 1 and 59 Hz, common average rereferencing, epoching from -1000 to +2000 ms around stimulus onset, and baseline correction using the window from -1000 to -300ms prior to stimulus onset. Further details on the removal of artifactual data can be found in Riddle et al., 2021.

We isolated our analyses of task data to two electrode clusters for each participant: a prefrontal channel cluster, and a posterior channel cluster. Based on our previous work, we hypothesized to find unique timescale dynamics in parietooccipital cortex that was distinct from prefrontal cortex. For the EEG data from Dataset 1, we averaged frontocentral electrodes (F1, Fz, F2, FC1, FCz, FC2) and parietooccipital electrodes (P3, P1, Pz, P2, P4, PO3, POz, PO4) that were identified using a hierarchical clustering analysis (Riddle et al., 2020). For Dataset 2, the channels used for the prefrontal cluster were Fz and the surrounding electrodes and POz and its surrounding electrodes for the posterior cluster. For each participant, each trial, and each electrode cluster, two windows of EEG data were extracted for analysis: one pre-stimulus window of time from -1000 to 0 ms, and a second post-stimulus window from +500 ms to +1500 ms. The post-stimulus window timing was selected to provide some buffer from the pre-stimulus window for estimating spectral activity. Additionally, this time window contained the majority of the stimulus-locked oscillatory results reported in Riddle et al., 2020 and Riddle et al., 2021.

### Outlier Removal and Statistical Analysis

We opted to analyze single-trial parameters, as opposed to model fitting exclusively on trial-averaged EEG data, in order to avoid the potential loss of data from whole participants/conditions where model fitting fails. However, the results of trial-averaged parameterization can be found in **Supplementary Notebook 02**. We first log-transformed raw 50-Crossing and aperiodic tau parameter values to enforce a normal distribution of the values. Data were then pooled across participants, conditions, and electrode clusters and assessed for outliers based on a 95% percentile exclusion criteria, where the upper and lower 2.5% of values were excluded from analysis.

Aperiodic PSD-Exponents, aperiodic ACF-Tau, and oscillatory PSD-Gaussian peak parameters from model fits with an R^2^ value lower than 0.5, or trials in which model fitting failed, were additionally removed from analysis. Aperiodic ACF-Tau parameters from model fitting conducted without a damped cosine have a systematically lower R^2^ value than those that include the damped cosine. This is due to the deliberate ignoring of the oscillatory effects on the ACF in an attempt to only capture the aperiodic “background” exponential decay function (**Figure 1D**). Therefore, in the cases where we excluded the damped cosine component as a hyperparameter of the model fitting procedure, we considered models with an R^2^ lower than 0.34 to be poor fits (selected to eliminate a similar proportion of the data as the 0.5 threshold for the alternative models). We include a sensitivity analysis in **Supplementary Notebook 01** for an assessment of how the model-inclusion R^2^ threshold choices of 0.5 and 0.34 might influence our statistical outcomes. These inclusion criteria were evaluated for both pre-stimulus time windows and post-stimulus time windows, and any trials wherein either window did not meet the criteria were dropped from analysis. Dataset 1 and 2 only included trials with correct task responses. After all of these data cleaning procedures were performed (separately for every neural metric), we retained an average of 79 trials per condition for use in our analyses.

Difference scores for each metric were calculated as the post-stimulus minus the pre-stimulus value, divided by the pre-stimulus value, and multiplied by 100 to derive a percent-change. The exception to this rule was the aperiodic-corrected oscillatory bandpower difference scores, which we calculated as an absolute difference between the pre- and post-stimulus values. We investigated the impact of task difficulty as the contrast high-difficulty conditions (R8, D2) from low-difficulty conditions (R4, D1), and task abstraction as the contrast of high-abstraction conditions (D1, D2) from low-abstraction conditions (R4, R8).

Resting-state data were submitted to the same parameterization and data cleaning procedures as described above for task data. The resulting set of parameters estimated from sequential 1-second windows of the continuous data were randomly shuffled. A subset of 79 “trials” were randomly selected from these shuffled datapoints, with the goal of matching the average number of task epochs we extracted for each task dataset (for each condition). We then computed the median. This procedure was repeated 100 times for each participant, and the set of 100 median values were averaged to give one resting-state datapoint per neural metric, per participant. We then compared these resting-state averages for each neural metric, and each participant, to those computed for the pre-stimulus and post-stimulus time windows of the task data. These data are plotted in **Figure 2** for the 50-Crossing metric, and in **Figure S1G**.

**Figure 2:**
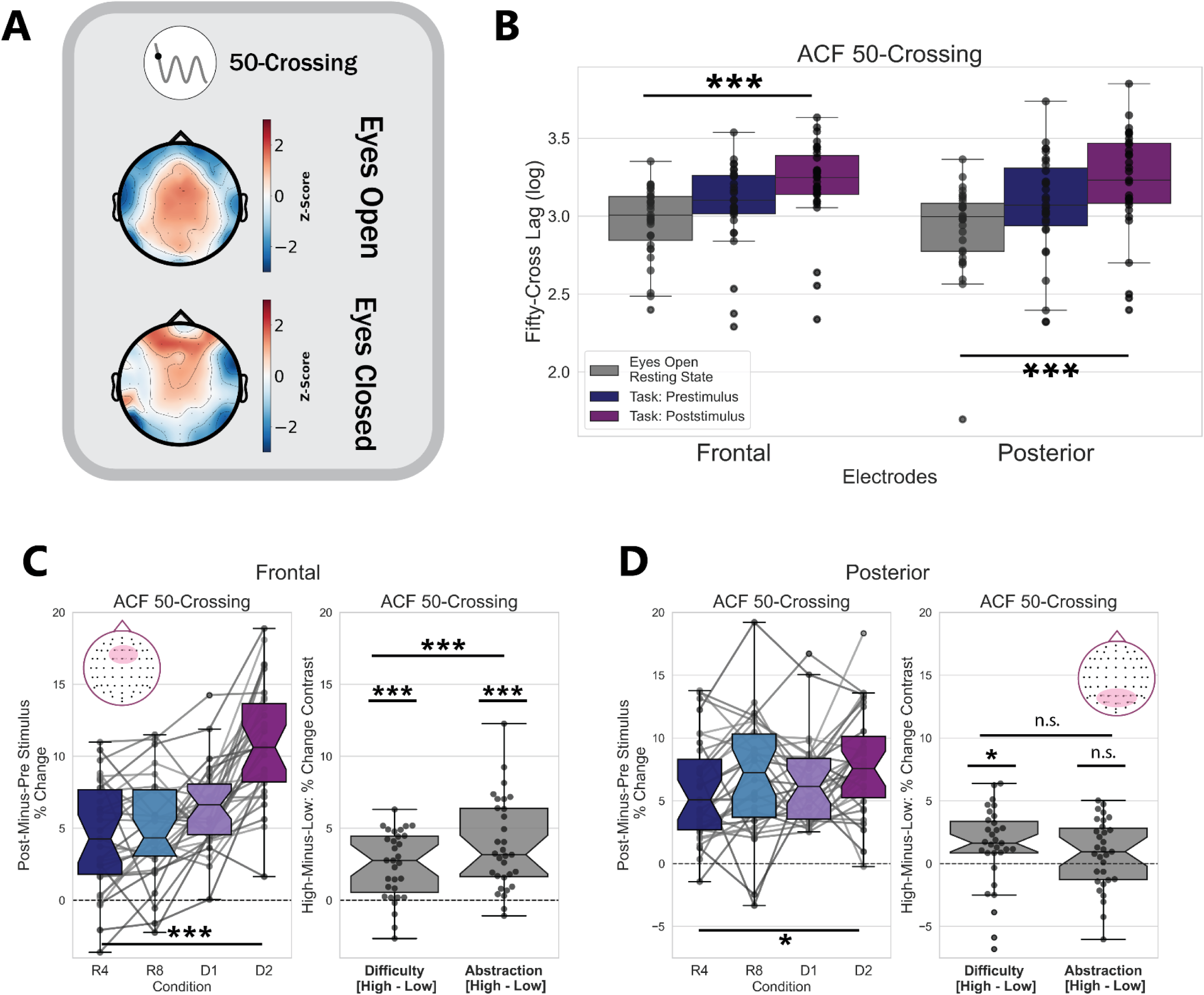
Traditional measures of neural timescales show timescale dilation between eyes-open resting state, pre-stimulus task periods, and post-stimulus task periods. A) Distribution of neural timescales over electrodes during eyes-open and eyes-closed resting state, from Dataset 1. B) Eyes-open resting state exhibits the fastest timescales, compared to pre- and post-stimulus windows, as measured by the 50-Crossing metric. Each point on the boxplot is the trial-averaged data for one participant, for each region of interest. C) (left) Trial-averaged logged 50-Crossing values, sampled from raw ACFs, lengthen pre-to-post stimulus in all conditions, with a main effect of both difficulty and abstraction, as well as an interaction between these (right). The 50-Crossing mixes both aperiodic and oscillatory activity. D) Same as C, but for the posterior channel cluster. The 50-Crossing here shows a significant main effect of condition, which seems to be driven by timescales lengthening selectively during high-difficulty conditions.

To assess the effect of condition on stimulus-evoked aperiodic and oscillatory activity, we ran repeated-measures analysis of variance (ANOVA) statistical tests on the trial-averaged tau aperiodic parameter, exponent aperiodic parameter, 50-Crossing, and oscillatory bandpower difference scores. We additionally tested the effects of task abstraction and task difficulty by conducting two-way repeated measured ANOVA tests, with trials averaged according to their respective levels of high/low abstraction/difficulty. We statistically tested the presence of FOOOF-detected oscillatory Gaussian peaks for alpha and theta bands, as they differed between the pre-stimulus and post-stimulus time windows, using McNemar’s test. This test is similar to a chi-squared test for independence, but it allows for observations to be non-independent. Lastly, we conducted a post hoc one-sample T-test to evaluate whether the change between pre-and-post stimulus of each metric was statistically significantly greater than zero. This procedure was done for each region of interest and data were collapsed across condition.

We additionally submitted these data to a series of linear mixed effects (LME) models, with the goal of examining the unique variance explained by each neural metric for single-trial reaction times. The full model (Model 1) we used as a point of comparison was formulated as:

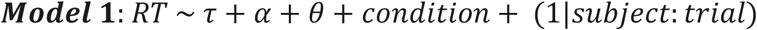

Where *RT* is the reaction time on every trial, and fixed effects terms 𝜏, 𝛼, and 𝜃 are the z-score normalized changes from baseline for the ACF-Tau (fit without oscillations), PSD-Alpha bandpower, and PSD-Theta bandpower for each trial. We compared this Model 1 to a model that only includes the 50-Crossing metric (Model 2); though we suspect that the 50-Crossing metric blends the contributions of aperiodic and periodic components to the ACF, we wished to examine whether it captured any variance in the reaction time data over and above that of Model 1. This Model 2 was formulated as:

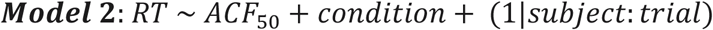

We additionally sought to test the hypothesis that the condition label alone could fully account for single-trial reaction time. As such, we compared Model 1 and Model 2 to a reduced model, Model 3, containing only a fixed term for condition:

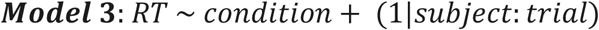

Model comparisons were performed using a likelihood ratio test for nested models (43), and Akaike Information Criterion (AIC) (44) for non-nested models. All LME-related procedures were conducted using the lme4 package (45) in RStudio (46). Each of these procedures were performed once for each channel cluster (Prefrontal and Posterior), and separately for Dataset 1 and Dataset 2.

### Replication Procedures

The data processing, analysis, and outlier removal pipeline was developed and finalized on Dataset 1 before application to Dataset 2. Statistical results for Dataset 2 are reported below, but all Dataset 2 figures can be found in **Figure S3 and S4**.

### Results: Dataset 1

#### Task Timescales are Longer than Resting-State Timescales

We quantified neural timescales using the traditional 50-Crossing measure on 1-second time windows of eyes-open resting state data (see Methods for details on resting-state analyses). We then compared these timescales to those found during the pre-stimulus and post-stimulus windows of task data, trial-averaged for each participant and electrode cluster. Submitting these data to a one-way rmANOVA revealed a main effect of task state on neural timescales for both the prefrontal (F(2, 60) = 25.38, p < 2x10^-8^, 𝜂^2^𝐺 = 0.136) and posterior (F(2, 60) = 16.99, p < 2x10^-6^, 𝜂^2^𝐺 = 0.122) channel clusters (**Figure 2B**).

Overall, neural timescales measured with a traditional metric— conflating aperiodic and oscillatory activity— show that timescales lengthen during pre-stimulus task states and lengthen more during post-stimulus task states, compared to eyes-open resting state. These results suggest that the integration demands of task states, in general, are reflected in longer windows of neural integration.

### Traditional Measurements Show Prefrontal Timescales Lengthening with Difficulty and Abstraction

We hypothesized that the 50-Crossing metric would show the greatest increase (lengthening) in the high-abstraction conditions, as these conditions should drive the greatest demand on integration for context-dependent stimulus processing.

The 50-Crossing metric lengthened pre-to-post stimulus, with the percent-change showing a significant difference from zero for both the prefrontal (*t*(29)=13.5, p < 2x10^-14^, *d* = 2.43) and posterior (*t*(29)=11.5, p < 8x10^-13^, *d* = 2.07) clusters. Both the prefrontal and posterior 50-Crossing did vary significantly with condition (prefrontal F(3, 30) = 37.13, p < 2x10^-15^, 𝜂^2^𝐺 = 0.341; posterior F(3, 30) = 3.11, p = 0.039, 𝜂^2^𝐺 = 0.036 after Greenhouse-Geisser correction for sphericity) in a one-way rmANOVA (**Figure 2C-D**).

A two-way rmANOVA of stimulus-locked changes in the 50-Crossing metric from prefrontal electrodes revealed a main effect of task abstraction (F(1, 30) = 46.10, p < 2x10^-7^, 𝜂^2^𝐺 = 0.237), a main effect of task difficulty (F(1, 30) = 31.78, p < 4x10^-6^, 𝜂^2^𝐺 = 0.109), and an interaction between these (F(1, 30) = 25.21, p < 3x10^-5^, 𝜂^2^𝐺 = 0.078). We found an additional main effect of task difficulty on the stimulus-locked lengthening in the posterior 50-Crossing metric (F(1, 30) = 5.03, p = 0.032, 𝜂^2^𝐺 = 0.026), with no main effect of task abstraction (F(1, 30) = 2.57, p =0.119, 𝜂^2^𝐺 = 0.009).

Overall, these findings support our hypothesis that neural timescales, particularly measured from prefrontal electrodes, lengthen during cognitive task abstraction.

### Prefrontal Theta Oscillations Emerge and Posterior Alpha Oscillations Disappear with Stimulus Onset

How might oscillatory signals become faster or slower with stimulus onset? Neural oscillations might change in terms of their power or center frequency, and over time may emerge or disappear altogether. We therefore began with inspecting the overall commonality of oscillatory peaks in the pre-and-post stimulus time windows by examining how many peaks our FOOOF models detected. In order to test the effect of time window on PSD Gaussian Peak presence, we pooled all trials with a successful PSD model fit, and with R^2^ values higher than 0.5 in both pre-stimulus and post-stimulus time windows, from all conditions and all participants.

For each trial, and each time window, we indicated whether the FOOOF fitting algorithm detected any oscillatory peak in the theta and alpha bands (1 for peak(s) detected, 0 for no peak detected). We limited the scope of our statistical analyses to the theta oscillatory peaks detected in the prefrontal electrode cluster, and alpha oscillatory peaks detected in the posterior electrode cluster, due to these sites being the canonical loci of theta/alpha oscillations during cognitive control (47,48). However, oscillatory peaks from the other electrode clusters are plotted in **Figure S2A**.

Notably, PSD Gaussian Peaks were not modeled in all or even the majority of trials, during either the pre-stimulus or post-stimulus windows. Prefrontal theta oscillatory peaks were detected in greater numbers (47% of trials) after stimulus onset than before stimulus onset (39% of trials) (**Figure S2B**). The opposite pattern held for posterior alpha oscillatory peaks; alpha oscillations were found in greater numbers pre-stimulus (53% of trials) compared to post-stimulus (42%). We submitted these data to McNemar chi-square tests for paired data in order to test whether trial counts of oscillatory peak presence and non-presence statistically differ pre-to-post stimulus. Both prefrontal theta (𝑋^2^(1, N=10,457) = 162, p < 5x10^-37^) and posterior alpha oscillatory peak presence (𝑋^2^(1, N=10,505) = 25.5, p < 5x10^-7^) showed a significant effect of time window.

These results suggest that one factor contributing to the slowing of neural activity post-stimulus may be the shift in oscillatory peak presence from predominantly faster (8-12 Hz alpha) to predominantly slower (4-8 theta) oscillations. Moreover, though these results replicate a classic finding of prefrontal theta power increase and concurrent posterior alpha power decrease in response to attended stimuli (47), they suggest that oscillatory peak presence may not be a constant of every trial, but rather a PSD feature emerging above threshold only for a subset of trials.

### Oscillatory Power Changes with Difficulty and Abstraction

In addition to prefrontal theta and posterior alpha oscillatory peak presence, we quantified stimulus-locked changes in oscillatory bandpower. We opted not to use the FOOOF-fit PSD Gaussian Peak parameters for this analysis, because they were missing for more than half of all trials. This meant that many participants/conditions were missing data, and statistical tests with ANOVAs were not possible. Instead, we calculated aperiodic-corrected oscillatory bandpower for every trial and time window. This method of quantifying oscillatory power assumes that trials with low-powered or no oscillations will have an aperiodic-adjusted bandpower estimate near zero. We then calculated the bandpower changes from pre-stimulus to post-stimulus as an absolute change, trial-averaged these within condition and within participant, and submitted them to our statistical tests.

In a post-hoc one-sample t-test, we found that prefrontal PSD-Theta power significantly increased from zero (*t*(29) = 9.5, p < 8x10^-11^, *d* = 1.70), while posterior PSD-Alpha bandpower decreased significantly (*t*(29) = -10.5, p < 7x10^-12^, *d* = 1.89). We submitted the trial-averaged, raw difference scores of oscillatory power for every condition to a one-way rmANOVA (**Figure 3**). We found a main effect of task condition on pre-to-post stimulus changes in prefrontal theta aperiodic-adjusted bandpower (F(3, 30) = 5.46, p = 0.002, 𝜂^2^𝐺 = 0.101), where prefrontal theta bandpower increased post-stimulus, on average, relative to pre-stimulus power. Likewise, we found a main effect of task condition on stimulus-locked changes in posterior alpha bandpower (F(3, 30) = 19.13, p < 2x10^-9^, 𝜂^2^𝐺 = 0.219), with alpha power decreasing the most in the high difficulty/high abstraction condition (D2).

**Figure 3:**
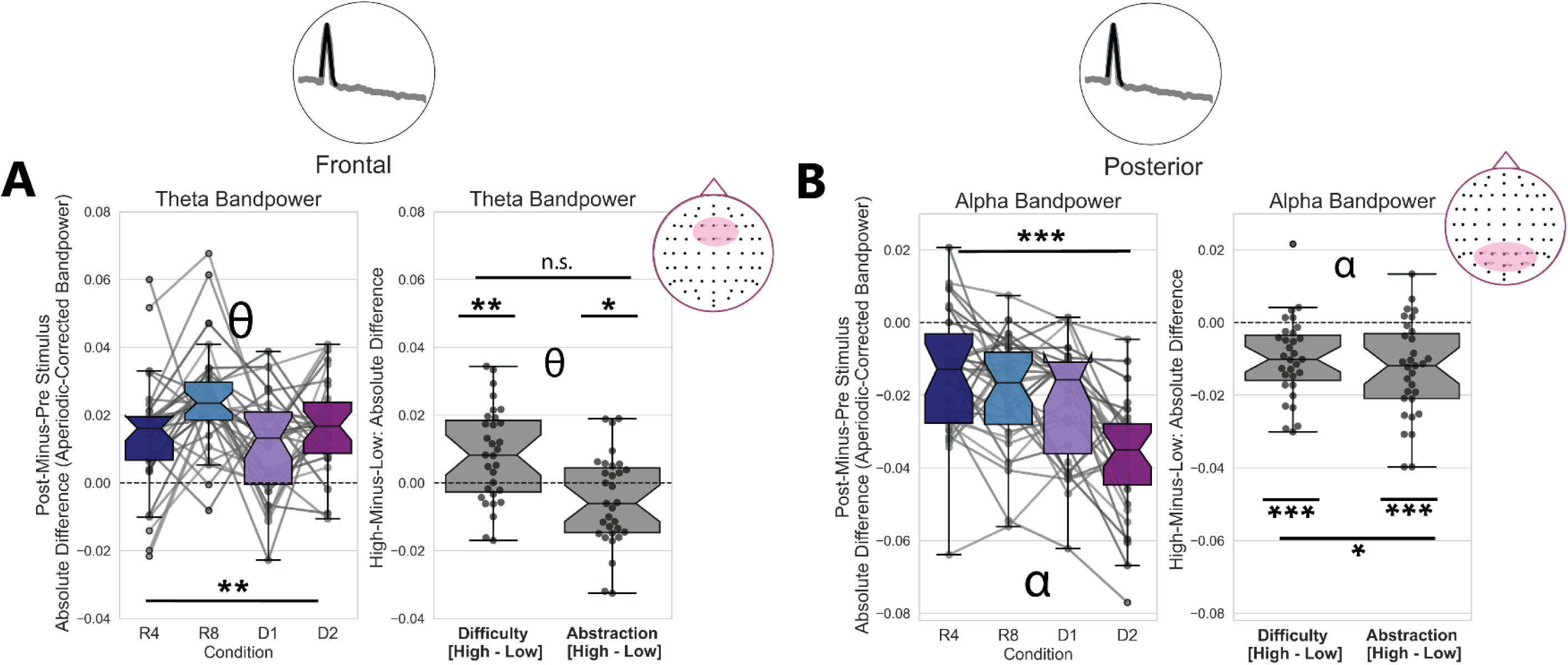
Aperiodic-Corrected Theta Oscillatory Power Increases with Task Difficulty, with Concurrent Decrease in Alpha Oscillatory Power. A) Frontal theta aperiodic-corrected bandpower (4-8 Hz) increases pre-to-post stimulus, with a main effect of Difficulty revealed by a two-way repeated-measures ANOVA. B) Posterior alpha aperiodic-corrected bandpower (8-12 Hz) decreases in the post-stimulus time window, relative to the pre-stimulus window. This alpha bandpower decrease showed a main effect of both abstraction and difficulty, as well as an interaction between these.

To further investigate whether changes in prefrontal theta oscillatory bandpower were driven selectively by either task difficulty or abstraction, we ran a two-way rmANOVA. We found a main effect of task difficulty (F(1, 30) = 11.99, p = 0.0016, 𝜂^2^𝐺 = 0.072), a main effect of task abstraction (F(1, 30) = 4.65, p = 0.039, 𝜂^2^𝐺 = 0.028), but no interaction between these (F(1, 30) = 0.82, p = 0.37, 𝜂^2^𝐺 = 0.007). We additionally found a main effect of task difficulty (F(1, 30) = 24.6, p < 3x10^-5^, 𝜂^2^𝐺 = 0.086), a main effect of task abstraction (F(1, 30) = 28.29, p < 1x10^-5^, 𝜂^2^𝐺 = 0.137) and an interaction between difficulty and abstraction (F(1, 30) = 5.08, p = 0.03, 𝜂^2^𝐺 = 0.023) on stimulus-locked changes in posterior alpha aperiodic-adjusted bandpower. These statistical results are reported in **Table 1**.

Altogether, these results suggest oscillatory neural timescales lengthen with task abstraction and difficulty. Specifically, prefrontal theta oscillatory power increases the most post-stimulus, relative to pre-stimulus, in the high-Difficulty conditions. Conversely, alpha oscillatory power in posterior electrodes decreases with stimulus onset, and decreases the most in the high-Abstraction/high-Difficulty condition (**Figure 3**). Thus, oscillatory activity slows post-stimulus via the joint emergence of, and increase of power in, oscillations in the theta band range alongside a concurrent decrease in alpha oscillation presence and power.

### Prefrontal Aperiodic Timescales Lengthen with Abstraction

We quantified changes in the aperiodic contribution to neural timescales first by using ACF parameterization to estimate Tau, or the inverse of the decay rate of the ACF, and secondly by PSD parameterization to estimate exponent of the PSD as a proxy for the aperiodic knee. We calculated the percent change in ACF-Tau and PSD-Exponent pre-to-post stimulus. These percentages were trial-averaged and then submitted to a one-way repeated-measured analysis of variance (rmANOVA) test for the effect of condition. These data were additionally submitted to a two-way rmANOVA to test for the pooled effects of task difficulty (high difficulty conditions were R8 and D2, low difficulty conditions were R4 and D1) and task abstraction (high abstraction conditions were D1 and D2, low abstraction conditions were R4 and R8). The results of these analyses, for both Dataset 1 and Dataset 2, are reported in **Table 1**.

We observed that the percent-change in ACF-Tau from baseline was significantly greater than zero in prefrontal (*t*(29)=9.34, p < 2x10^-10^, *d* = 1.67) and posterior (*t*(29)=10.6, p < 6x10^-12^, *d* = 1.90) sites. The percent-change in the prefrontal ACF-Tau parameter has a main effect of condition (F(3, 30) = 4.39, p = 0.006, 𝜂^2^𝐺 = 0.059), with a main effect of abstraction (F(1, 30) = 9.36, p = 0.005, 𝜂^2^𝐺 = 0.041), no main effect of difficulty (F(1, 30) = 0.50, p = 0.48, 𝜂^2^𝐺 = 0.003), and an interaction between difficulty and abstraction (F(1, 30) = 4.68, p = 0.038, 𝜂^2^𝐺 = 0.017). These data are plotted in **Figure 4**. Data for the posterior channel cluster did not show any significant main effects of condition or task abstraction/difficulty, and the statistical results of the rmANOVA for these data can be found in **Table 1**.

**Figure 4:**
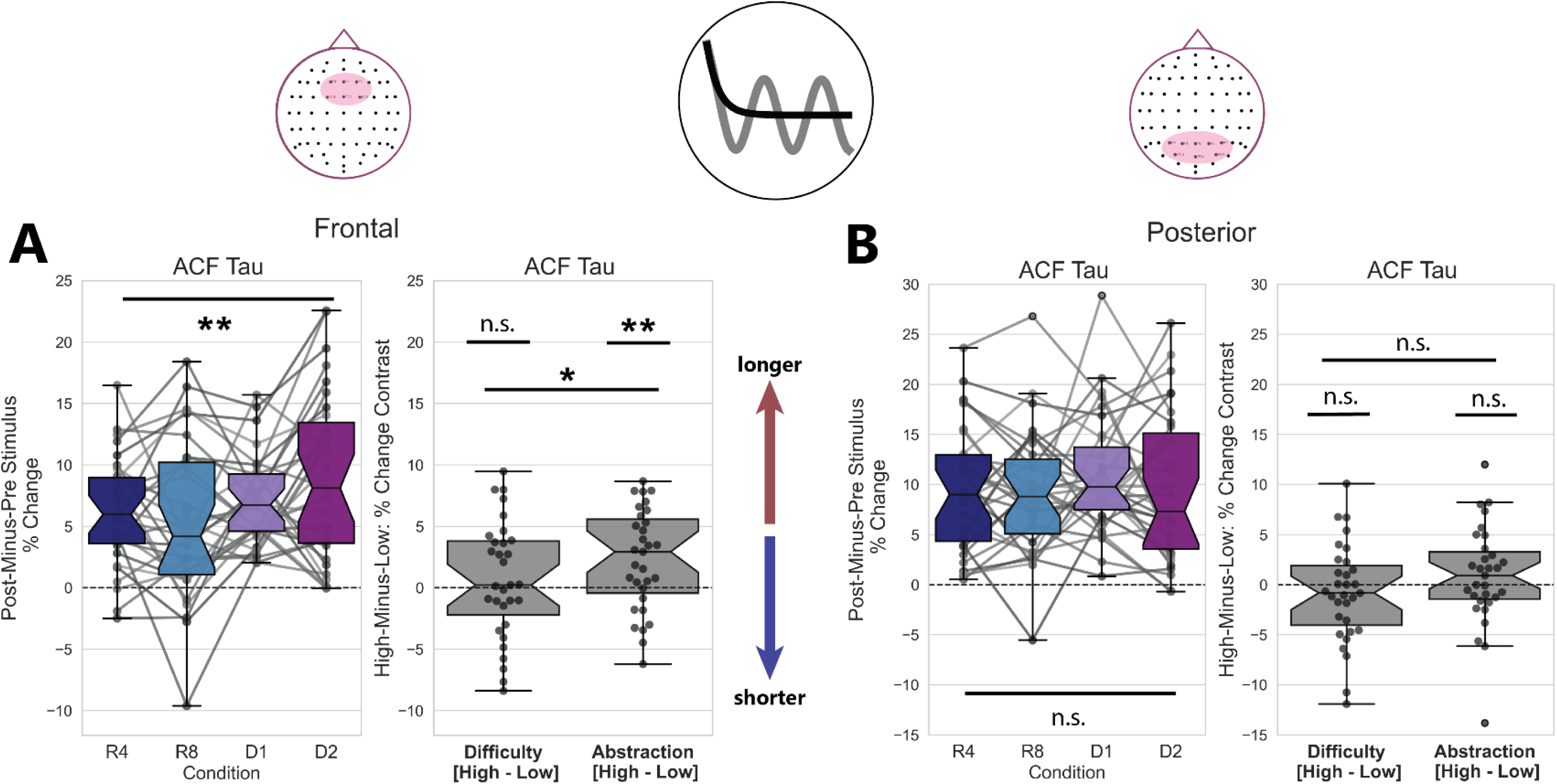
Aperiodic Timescales in Prefrontal Electrodes Lengthen with Abstraction. A) Timescales measured from trial-averaged logged ACF Tau parameters (left) lengthen the most in the high-Difficulty, high-Abstraction condition. ACF Tau lengthening (right) from pre-to-post stimulus shows a significant main effect of Abstraction, no main effect of Difficulty, and a significant interaction between Abstraction and Difficulty in a two-way repeated-measures ANOVA. B) Though they increase overall post-stimulus, the ACF Tau parameter measured at Posterior electrodes does not exhibit a condition-specific increase pre-to-post stimulus.

One-way rmANOVAs for prefrontal PSD-Exponent parameters (**Figure S2C-D**) revealed a trending effect of condition (F(3, 30) = 2.37, p = 0.076, 𝜂^2^𝐺 = 0.041). However, a post-hoc one-sample t-test revealed that the PSD-Exponent increases significantly from zero in both prefrontal (*t*(29)=22.5, p < 2x10^-20^, *d* = 4.05) and posterior (*t*(29)=10.3, p < 2x10^-11^, *d* = 1.85) sites, when data are collapsed across condition. Analyses of the prefrontal channel cluster suggest no main effect of task abstraction (F(1, 30) = 2.03, p = 0.165, 𝜂^2^𝐺 = 0.017), and a trending but not significant main effect of task difficulty (F(1, 30) = 3.88, p = 0.058, 𝜂^2^𝐺 = 0.019). We conducted the same set of statistical tests for the PSD-Exponent parameters derived from a cluster of posterior electrodes. The full results of this procedure can be viewed in **Table 1**; these did not show any main effects of condition, Abstraction, or Difficulty.

Overall, these findings suggest that aperiodic timescales measured at prefrontal sites exhibit lengthening post-stimulus, where such lengthening is best captured by measuring from the ACF. This lengthening is also condition-specific, with the highest-abstraction task contexts driving aperiodic timescales lengthening to the greatest degree.

### Neural Timescales Predict Reaction Times Independently of Condition

One possibility is that reaction time is not predicted by the slowing in neural timescales, but predicted wholly by the effects of the task condition. We thus used a series of LME models to examine the relationship between single-trial reaction times and each of our neural metrics. The models are formulated as follows (see Methods section for details):

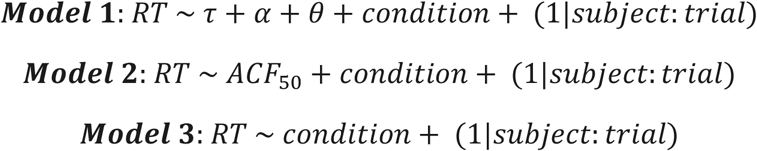

We first compared the full model, Model 1, to a reduced null Model 3, to assess whether our neural metrics increase the precision of the model more than the condition label alone. For prefrontal channels, this showed a significant effect of ACF-Tau, PSD-Alpha bandpower, and PSD-Theta bandpower (𝜒^2^(3) = 30.21, p < 2x10^-6^; Model 1 AIC < Model 3 AIC), as did the model trained on posterior-cluster metrics (𝜒^2^(3) = 35.34, p < 2x10^-7^; Model 1 AIC < Model 3 AIC). We additionally tested whether the 50-Crossing metric correlated with reaction time, independently of condition label, by comparing Model 2 with the null Model 3. These analyses revealed a main effect of 50-Crossing for single-trial reaction times for both the prefrontal cluster (𝜒^2^(1) = 34.28, p < 5x10^-9^; Model 2 AIC < Model 3 AIC) and the posterior cluster (𝜒^2^(1) = 5.88, p = 0.015; Model 2 AIC < Model 3 AIC).

We additionally asked whether the 50-Crossing metric, as a likely combination of the aperiodic and periodic components, is more informative for reaction time (Model 2) than measuring each of the metrices separately as in Model 1. We found a lower AIC for Model 2 (1402.88) than for Model 1 (1430.64) for the prefrontal channels, but not for the posterior channels (Model 2 AIC = 1313.85, Model 1 AIC = 1307.85).

Overall, these results suggest that the aperiodic and oscillatory contributions to neural timescales quantified in the present study capture some effects on single-trial reaction time that the task condition alone does not fully account for.

### Replication Results: Dataset 2

#### Replication of Traditional Neural Timescale Measure (50-Crossing) Slowing with Abstraction in Prefrontal Electrodes

Our 50-Crossing results were replicated in Dataset 2, with a significant change from zero in prefrontal (*t*(21) = 9.1, p < 4x10^-9^, *d* = 1.89) and posterior (*t*(21) = 8.2, p < 2x10^-8^, *d* = 1.71) sites. In the prefrontal electrode cluster for Dataset 2, we observed a main effect of condition (F(3, 22) = 19.77, p < 2x10^-7^, 𝜂^2^𝐺 = 0.180 after Greenhouse-Geisser correction for sphericity), and a main effect of both abstraction (F(1, 22) = 38.04, p < 4x10^-6^, 𝜂^2^𝐺 = 0.117) and difficulty (F(1, 22) = 15.05, p = 0.0008, 𝜂^2^𝐺 = 0.072) (**Figure S3C**). The findings for the 50-Crossing measured from posterior channels (**Figure S4C**) did not replicate those of Dataset 1; we found no significant main effect of condition, abstraction, or difficulty (**see Table 1**).

#### Replication of Prefrontal Theta Emergence and Posterior Alpha Disappearance with Stimulus Onset

We replicated Dataset 1’s oscillatory peak presence findings in Dataset 2: prefrontal theta peak presence increased post-stimulus compared to pre-stimulus (𝑋^2^(1, N=7,940) = 167.94, p < 3x10^-38^), with a concurrent decrease in posterior alpha peak presence (𝑋^2^(1, N=8,062) = 185, p < 5x10^-42^) (**Figure S3D-E**).

In a replication of the findings from Dataset 1 (**Figure 3A**), we found that PSD-Theta bandpower increases significantly (*t*(21)=8.8, p < 6x10^-9^, *d* = 1.84), while posterior PSD-Alpha bandpower decreases significantly from zero (*t*(21) = -8.2, p < 2x10^-8^, *d* = 1.72). We found that prefrontal theta and posterior alpha oscillatory bandpower showed a main effect of condition (theta: F(1, 22) = 3.79, p = 0.01, 𝜂^2^𝐺 = 0.097; alpha: F(1, 22) = 26.1, p < 4x10^-11^, 𝜂^2^𝐺 = 0.279). We additionally observed main effects of difficulty for both prefrontal theta bandpower and posterior alpha bandpower, with an additional main effect of abstraction for posterior alpha (see **Table 1**). The only finding that was not replicated in Dataset 2 was a lack of a significant effect for task abstraction for prefrontal theta bandpower.

#### Replication of Aperiodic Timescales Lengthening with Abstraction in Prefrontal Channels

The ACF-Tau fit *without* a oscillatory component (**Figure S3A**) showed a significant increase from zero in prefrontal (*t*(21)=17.8, p < 7x10^-15^, *d* = 3.73) and posterior (*t*(21)=13.1, p < 4x10^-12^, *d* = 2.74) sites. In a replication of Dataset 1, ACF-Tau also showed a main effect of condition for the prefrontal (F(3, 22) = 4.79, p = 0.009, 𝜂^2^𝐺 = 0.119 after Greenhouse-Geisser correction for sphericity), but not the posterior site (F(3, 22) = 0.35, p = 0.78, 𝜂^2^𝐺 = 0.007). The prefrontal site showed a main effect of abstraction (F(1, 22) = 7.47, p = 0.012, 𝜂^2^𝐺 = 0.081), no main effect of difficulty (F(1, 22) = 2.76, p = 0.11, 𝜂^2^𝐺 = 0.034), and a trending interaction between these (F(1, 22) = 3.24, p = 0.086, 𝜂^2^𝐺 = 0.012). For the posterior site (**Figure S4A**), there was no main effect of condition, abstraction, or difficulty (see **Table 1**).

In line with the main findings of Dataset 1, the stimulus-locked percent-change in PSD-Exponent is significantly greater than zero in both prefrontal (*t*(21)=12.7, p < 7x10^-12^, *d* = 2.65) and posterior (*t*(21)=10.1, p < 5x10^-10^, *d* = 2.12) sites. The PSD-Exponent (**Figure S3B and S4B**) did not vary significantly across condition, nor show any main effect of abstraction, and exhibited a trending main effect of difficulty for the prefrontal cluster (F(1, 22)=3.49, p = 0.07, 𝜂^2^𝐺 = 0.013) but not for the posterior cluster (see **Table 1**).

#### Replication of Neural Metrics, and Not Condition Label Alone, Predicting Single-Trial Reaction Times

Dataset 2 replicates the findings from Dataset 1. The comparison between Model 1 and the null Model 3 for the prefrontal channels showed a significant effect of the additional fixed parameters for ACF-Tau, PSD-Alpha, and PSD-Theta (𝜒^2^(3) = 26.11, p < 1x10^-5^; Model 1 AIC < Model 3 AIC), as did the posterior channels (𝜒^2^(3) = 18.73, p = 0.0003; Model 1 AIC < Model 3 AIC). The comparison between Model 2 and Model 3 revealed a main effect of the 50-Crossing on reaction times for the prefrontal (𝜒^2^(1) = 19.93, p < 9x10^-6^; Model 2 AIC < Model 3 AIC) and posterior channels (𝜒^2^(1) = 5.42, p = 0.02; Model 2 AIC < Model 3 AIC).

The comparison between our 50-Crossing Model 2 and the separate aperiodic/oscillatory parameter Model 1 suggests that, for prefrontal channels, Model 2 is more informative (AIC = 1203.74) than Model 1 (AIC = 1220.41). This pattern held for posterior channels (Model 2 AIC = 871.93; Model 1 AIC = 881.0

## Discussion

In the current study, we asked whether neural timescales exhibit task-evoked changes, and whether these changes vary with task abstraction. We found that neural timescales lengthen pre-to-post stimulus during a hierarchical cognitive control task. This held true for all neural metrics measured from scalp EEG in two independent datasets. We additionally found that timescale lengthening with task abstraction was specific to prefrontal aperiodic timescales, while oscillatory power showed sensitivity to task difficulty alone (in the case of prefrontal theta oscillations), or mixed sensitivity (posterior alpha oscillations).

We began our investigation by quantifying neural timescales using the traditional 50-Crossing measure, which samples from the raw ACF and therefore mixes aperiodic and oscillatory components of neural data. This introduces a major interpretational confound in that one cannot adjudicate between task-related changes in oscillatory features and task-related changes in non-oscillatory timescales. We therefore extended our analyses by separately modeling aperiodic timescales and oscillatory power. In doing so, we found that the ACF-Tau measured from prefrontal channels exhibited the greatest lengthening in the high-abstraction conditions. The PSD-Exponent showed lengthening in both frontal and posterior channels, though not in a condition-specific manner. PSD-Theta and Alpha oscillations exhibited increasing and decreasing aperiodic-corrected bandpower, respectively. While alpha band activity was sensitive both to task abstraction and task difficulty, theta band oscillations increased selectively in the high-difficulty conditions. That is, we found a tradeoff in power of a faster frequency oscillation, alpha, in favor of the increase in slower theta oscillations. This, in combination with the aperiodic results, indicates an overall shift in the neural activity from faster, higher-frequency pre-stimulus activity to slower activity after stimulus onset. The 50-Crossing metric demonstrated the greatest degree of timescale lengthening pre-to-post-stimulus with main effects of both abstraction and difficulty. That the 50-Crossing varied with both task abstraction and difficulty suggests that it is not capable of disambiguating the effects of the task on aperiodic and periodic activity, separately.

The limitations of the current study are primarily with the feasibility of modeling the aperiodic knee parameter from the PSD, as well as with measuring oscillations in the ACF. While the knee of the power spectrum is theoretically most directly related to the ACF-Tau, it is inappropriate to model the knee for spectra that are best fit as a line in semi-log space (36,49).

Here, our data showed heterogeneity in terms of whether the raw power spectra exhibited visible knees. We thus deferred to fitting a model without knees, and instead used the PSD-Exponent as a proxy (24,50). Though supplementary analyses suggest that our ACF-Tau findings might be replicated in the PSD-knee (**Supplementary Notebook 00**), further research is necessary to examine whether the knee of the power spectrum exhibits task-evoked changes in timescales. On a similar note, modeling the oscillatory effects on the ACF presents a unique set of challenges.

While the center frequencies of oscillations are theoretically recoverable from their effects on the ACF, the presence of multiple oscillations (such as the simultaneous theta and alpha oscillations observed in the current study) show up in the ACF as overlapping, nested oscillations. We therefore suggest that, for clean separation of the distinct oscillatory signals in the data, modeling in the PSD space may be a simpler and more reliable procedure than modeling the ACF. The ACF parameterization method presented here was only able to capture the effects of the dominant oscillation, ignoring any additional oscillations that were overlaid on it in the raw ACF.

Lastly, delta oscillations were outside of our window of observation given the brief time windows we analyzed from the task; with time windows of 1 second, we could not reliably fit a Gaussian peak on either side of a center frequency of 2Hz in the power spectrum. While previous studies found coupling of low-frequency delta and theta oscillations with motor-related beta oscillatory power (28,51,52) and low-frequency coupling with posterior alpha power (53), these findings might be confounded with changes in the power of low-frequency aperiodic activity such as the lengthening of timescales found in the present study (49,54). Conversely, recent work suggests that genuine delta oscillations exhibit coupling within the lateral prefrontal-parietal network while participants are performing the same task analyzed here (55). Thus, further research is necessary to dissociate task-sensitive dynamics of neural timescales from those that are specifically attributable to narrow-band delta oscillatory activity.

Though neural timescales have been posed as a fundamental feature of cortical organization (5,7,12,56), and this has been recapitulated in a multitude of imaging modalities (8), species (57,58), and brain states (33,59,60), there is a paucity of work examining neural timescales as a signature of functional re-organization during task states (13). Studies that quantify neural timescales during task performance are primarily using invasive EEG in non-human primates (13,61–64), and do not often investigate timescales as a dynamic feature of cortical regions (e.g., by measuring changes in timescales across different time windows of the task)—though see Golesorkhi et al., 2021 and Miller & Constantinidis, 2024 for notable exceptions.

The studies that do investigate task-evoked changes in neural timescales report a mix of results, with some showing that timescales are dynamic during a task (13,24), while others conclude that timescales are task invariant (26). Moreover, neural timescales have not been systematically studied in the context of task abstraction, or context-specific task processing. This presents a compelling gap in the literature, since it has been well established that the long-timescale regions of the prefrontal cortex are also those that are most engaged in abstraction during hierarchical cognitive control. That is, caudal regions of the frontal cortex track the planning and execution of concrete motor action, whereas more rostral regions track progressively higher-order, abstract task goals and context (20,29,66–72). Indeed, the dominant model accounting for the hierarchy of “intrinsic” neural timescales is that it reflects the progressive integration of information across the anatomical hierarchy of cortex (3,17,73,74)—a purported mechanism of task abstraction (68). The closest analogous research on timescales focuses on abstract linguistic and semantic processing, and is mostly conducted using fMRI. This literature broadly suggests that continuous low-level sensory inputs received by fast-timescale brain regions are progressively integrated into abstract and context-sensitive representations by higher-order, longer-timescale brain regions (18,75–85).

One additional factor that may add to the field’s ambiguous findings is the variety of ways that neural timescales are measured (17). Though the ACF is widely used to quantify timescales, the particular method for extracting “the” timescale from the ACF vary (6,33,86). Neural signals are composed of both rhythmic and non-rhythmic activity (2,30,36), and thus, the question of the timescale of a neural signal might be aptly described either in terms of aperiodic activity, oscillatory activity, or some combination of both. Commonly used N-Crossing methods for measuring the timescale, such as the 50-Crossing used in this study, likely capture some combined contribution of oscillatory and non-oscillatory dynamics in the underlying timeseries. Here, we employ several modeling approaches for estimating the aperiodic, oscillatory, and combined timescales of EEG data— each of which have been demonstrated in previous studies (6,16,24,33), but never compared all together.

Neural timescales are emerging as an important measure of cognitive processing as it progresses continuously and over multiple simultaneous timescales. That is, rather than reflecting an intrinsic, stable property of a given region of cortex, neural timescales may instead reflect the range of temporal receptive capacities of the region. Neural timescales therefore present a potentially unique lens through which to examine how functional dynamics within a task state interact with the large-scale anatomical gradients of the brain.

## Code Accessibility

The code used for main analyses and plots are publicly available on the Open Science Framework at https://osf.io/39kn7/?view_only=2d380f77a38f4c6d8e563356c8813254. A series of Jupyter notebooks outlining extended analyses can also be found at this link.

## Software Accessibility

Autocorrelation Function parameterization was performed using the timescales-methods toolbox, publicly available here: https://voyteklab.com/timescale-methods/.

Spectral parameterization was performed using the toolbox here: https://fooof-tools.github.io/fooof/.

## Acknowledgements

Support: e.g., NIH National Institute of General Medical Sciences grant R01GM134363-01 to B.V. R00MH126161 to J.R. NSF GRFP to D.C.

We thank Mark D’Esposito for his generously allowing us to re-analyze data collected in his lab, and Mattia Pagnotta for facilitating access to the data. We additionally thank Quirine van Engen for her support and contributions to the methodology of this research.

## Author contributions

D.C. coded/executed analyses and statistics. D.C., J.R., and B.V. developed analysis approach. J.R. developed the task protocol and published existing work related to these data. R.H. developed and consulted on the Auto-Correlation Function parameterization toolbox. D.C. and J.R. collected one dataset at University of California, Berkeley, and J.R. and F.F. collected a second dataset at University of North Carolina at Chapel Hill. D.C., J.R., F.F., and B.V. wrote and edited this paper.

## Supplementary Materials

Supplementary figures include plots of oscillatory and PSD-Exponent results of Dataset 1, and the results for all neural metrics in Dataset 2. Extended analyses can be found in the form of Jupyter notebooks hosted here: https://osf.io/39kn7/?view_only=2d380f77a38f4c6d8e563356c8813254.

---

**Figure S1:**
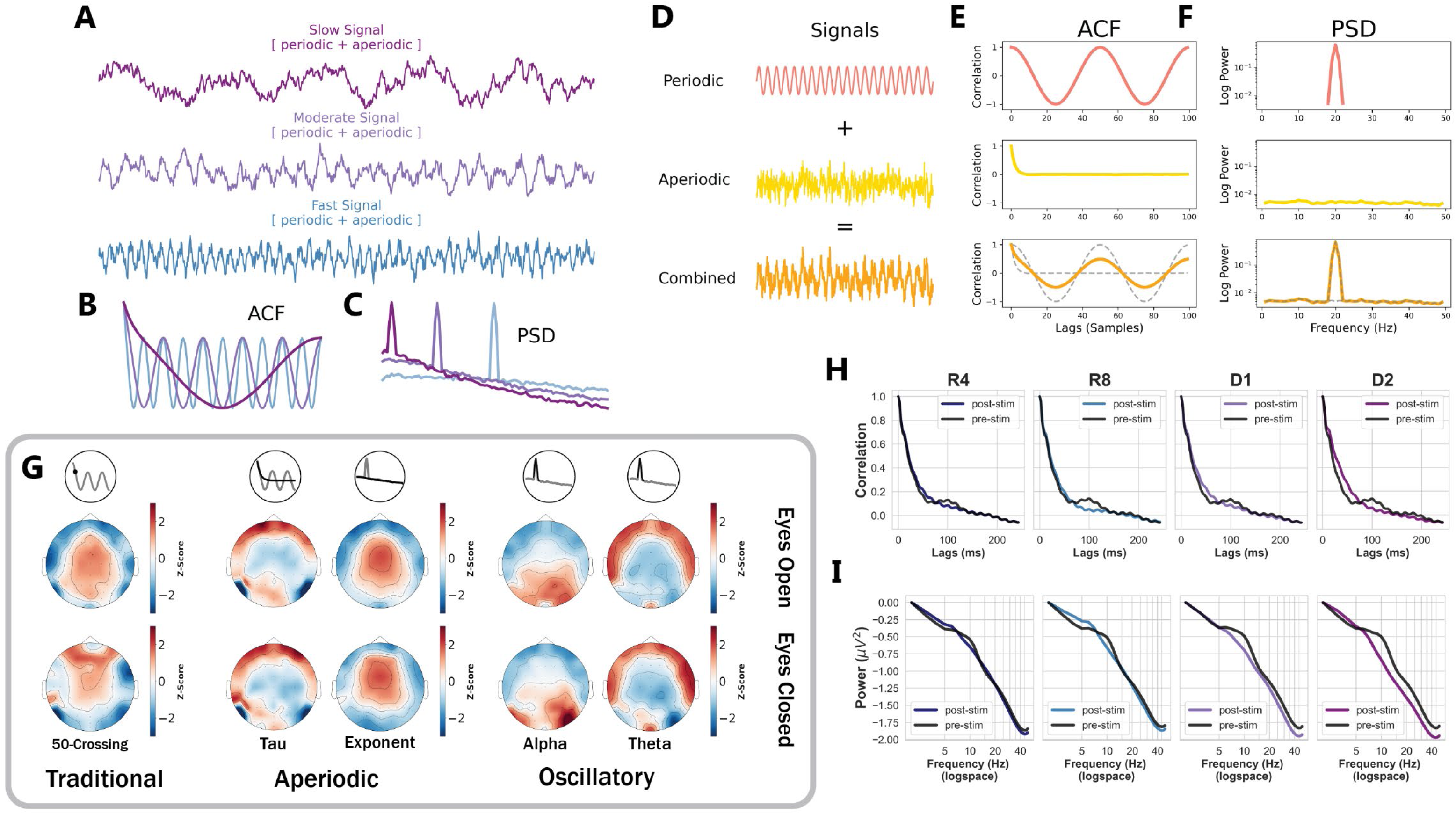
Neuronal timescales can be measured in either the Autocorrelation Function (ACF) or Power Spectral Density (PSD) space: A) Simulated signals with rhythmic (oscillatory) and non-rhythmic (aperiodic) components, generated to show 3 different “levels” of low-to-high frequency activity. Neural signals can vary in terms of their overall composition of stable, low-frequency, and dynamic, high-frequency signals. B) An autocorrelation function can be conceptualized as describing the rate of decay in the self-similarity of a timeseries signal. Signals that are “faster,” or closer to white noise, result in ACFs that decay quickly. Conversely, signals that are “slower,” or closer to pink/brown noise, result in ACFs that take more lags to decay (extending out further into the x-axis). Oscillations in the time domain of a signal manifest as oscillatory-like activity in the ACF, and likewise reflect the frequency of the oscillation. C) The power spectral density plots of the three simulated signals in A. Signals closer to white noise manifest in the PSD as flatter spectra, while the slowest signal yields a steeper spectrum. Oscillations appear as peaks over and above the “background” of the aperiodic activity. The knee of the aperiodic component of the power spectrum can be used to mathematically approximate the decay rate of the ACF. D) Simulated signals: a 20 Hz sine wave (red, top) combined with aperiodic activity (yellow, middle) produces a signal (orange, bottom) that mimics neural data in its combination of rhythmic and non-rhythmic signals. E) and F) The respective autocorrelation functions (ACF) and power spectral densities (PSD) resulting from transformations of the corresponding signals in A. While a pure sine wave in the time domain appears similarly rhythmic in the ACF space (E, top), it appears as a peak in the PSD space centered at the oscillatory frequency (F, top). The combination of oscillatory and aperiodic signals produces ACFs and PSDs (E and F, bottom) that resemble those derived from neural data. G) Distribution of each neural metric (from left to right: 50-Crossing, ACF-Tau, PSD-Exponent, PSD-Alpha bandpower, and PSD-Theta bandpower) over electrodes during eyes-open and eyes-closed resting state, from Dataset 1. H) Subject and trial-averaged raw ACFs for the pre- and post-stimulus windows, for each condition. The high-abstraction, high-difficulty D2 condition shows visibly slower timescales that manifest as longer decays in the ACF. I) Same as H, but showing raw PSDs plotted with offset-correction to align all spectra to zero. PSD knees at lower frequencies correspond to slower timescales. Averaging across subjects and trials may also distort oscillatory peaks in the underlying data, necessitating parameterization at a more granular level.

**Figure S2:**
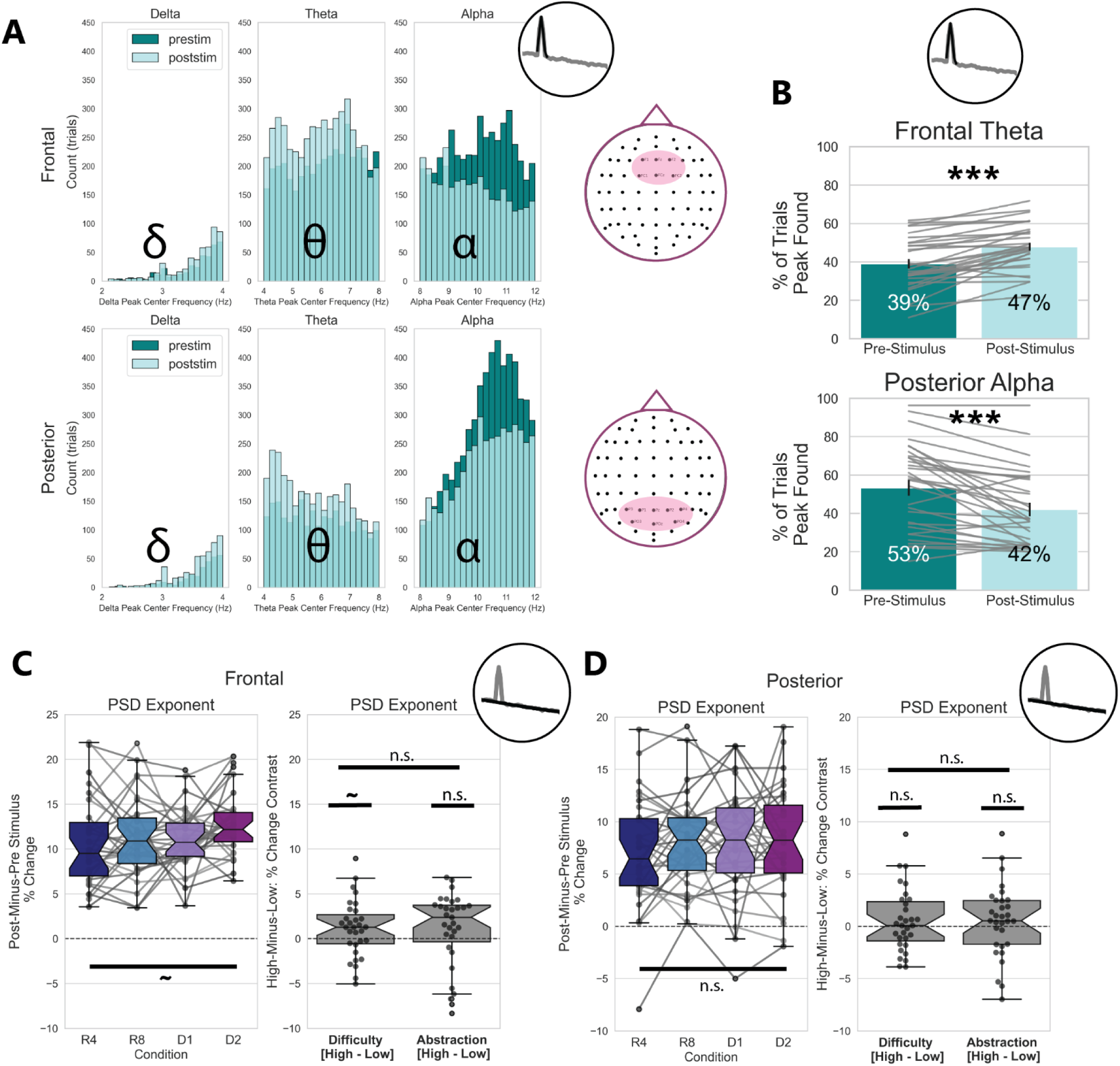
Oscillatory Peak Presence Differs between Pre-and-Post Stimulus Time Windows. A) Histograms of oscillatory peaks detected from PSD Oscillatory Peak model fitting, pooled across all subjects, all conditions, and all trials. Oscillatory peaks detected in the theta range (4-8 Hz) in Frontal Channels (top) are more numerous after stimulus onset, while oscillatory peaks detected in the alpha range (8-12 Hz) in Posterior Channels (bottom) decrease after stimulus onset. Model fits of delta oscillatory peaks at the lower bound edge of the power spectrum (2 Hz) are considered artifactual and were removed from plotting here. B) Same as A, but plotted in terms of the percent of all trials. PSD parameterization only successfully detected oscillatory peaks in a maximum of 53% of trials (pre-stimulus Alpha peaks), and the numbers of detected peaks shifts pre-to-post stimulus for both theta and alpha peaks. A McNemar test for significance revealed a significant effect of time window and oscillation type on oscillatory peak presence. C) and D) The trial-averaged aperiodic measure PSD-Exponent from Frontal and Posterior channels did not vary significantly across conditions, though it was trending (p = 0.076) in Frontal channels. They did not show a main effect of either Difficulty or Abstraction.

**Figure S3:**
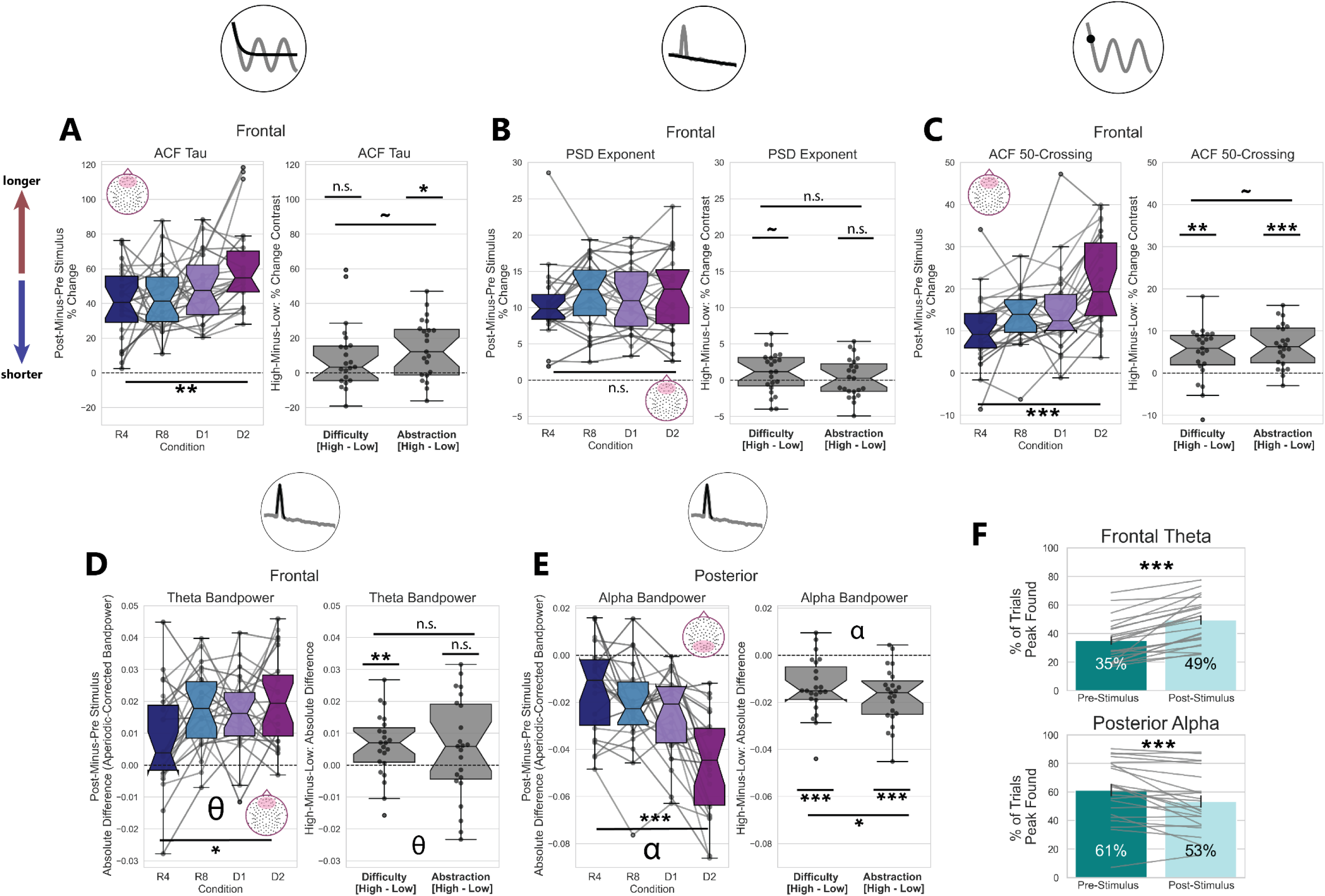
Aperiodic and oscillatory timescales lengthening from pre-to-post stimulus replicated in independent dataset. A) The ACF Tau parameter measured at Frontal electrodes in Dataset 2 exhibits a condition-specific increase pre-to-post stimulus. This finding replicated for the main effect of Abstraction, but not for the Abstraction and Difficulty interaction. B) The Frontal PSD Exponent did increase post-stimulus relative to pre-stimulus, but this increase did not vary significantly across conditions. C) The Frontal 50-Crossing measure from the ACF exhibited a significant increase, with a main effect of condition, Abstraction, and Difficulty. These data replicate the previous dataset, with the exception of finding only a trending interaction between Abstraction and Difficulty. D) and E) Frontal aperiodic-corrected theta bandpower replicated the previously observed main effect of condition, and a main effect of Difficulty, but not the main effect of Abstraction. The previous findings of posterior aperiodic-corrected alpha bandpower decrease, with main effects of condition, Abstraction, and Difficulty, as well as an interaction between Abstraction and Difficulty were replicated. F) In line with findings from Dataset 1, the number of theta oscillatory peaks detected by PSD parameterization decreased after stimulus onset; the reverse was true for alpha oscillatory peaks.

**Figure S4:**
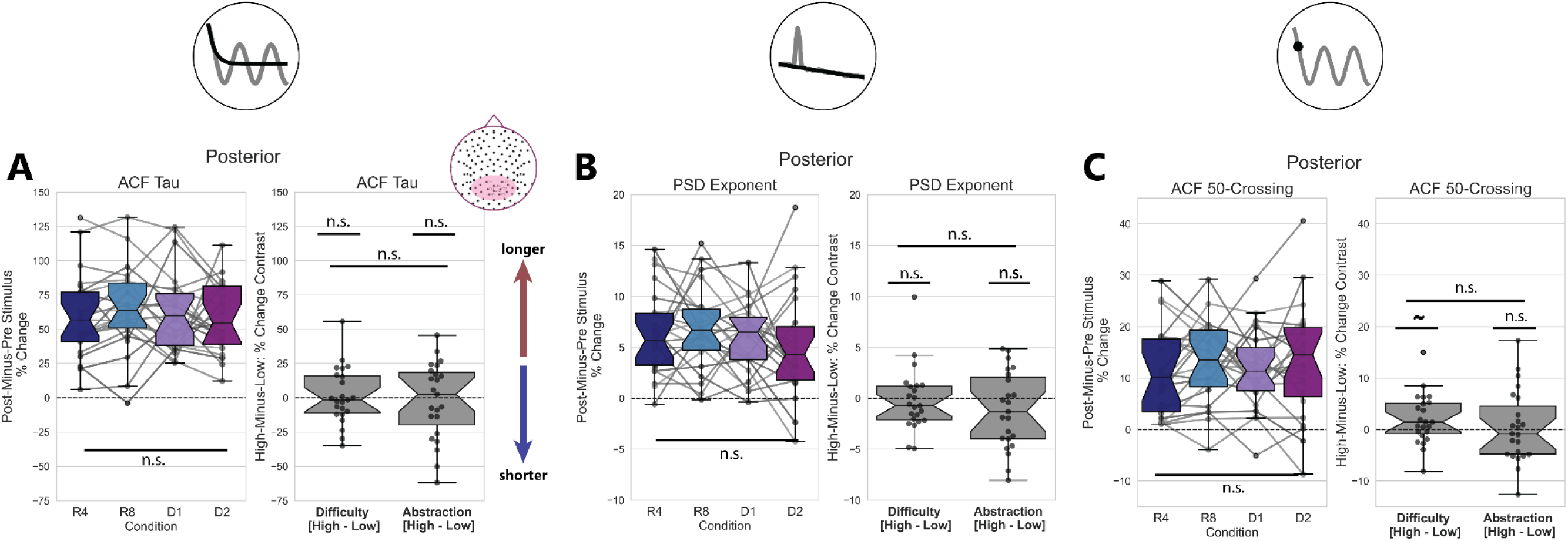
Aperiodic and oscillatory timescales lengthening from pre-to-post stimulus replicated in independent dataset. A) Though it increases overall post-stimulus, the ACF Tau parameter measured at Posterior electrodes does not exhibit a condition-specific increase pre-to-post stimulus. This finding is similar to that of Dataset 1. B) The Posterior PSD Exponent did increase post-stimulus relative to pre-stimulus, but this increase did not vary significantly across conditions. C) The Posterior 50-Crossing measure from the ACFs of Dataset 2 did not show a main effect of condition, unlike the findings of Dataset 1. These data did exhibit a trending but non-significant effect of Difficulty.

